# Variational calculus approach to Zernike polynomials with application to fluorescence correlation spectroscopy

**DOI:** 10.1101/2024.04.06.588390

**Authors:** Ivan Gligonov, Jörg Enderlein

## Abstract

Zernike polynomials are a sequence of orthogonal polynomials that play a crucial role in optics and in particular in modeling microscopy systems. Introduced by Frits Zernike in 1934, they are particularly useful in expressing wavefront aberrations and thus imperfections of imaging systems. However, their origin and properties are rarely discussed and proven. Here, we present a novel approach to Zernike polynomials using variational calculus, and apply them to describe aberrations in fluorescence microscopy. In particular, we model the impact of various optical aberrations on the performance of one-photon and two-photon excitation fluorescence microscopy.

**SIGNIFICANCE:** This manuscript explores the mathematical derivation of Zernike polynomials and highlights their critical role in describing optical wavefronts and aberrations, particularly in the domain of optical microscopy. Special emphasis is placed on their utility in simulating the effects of aberrations on one-photon and two-photon excitation fluorescence correlation spectroscopy.

## INTRODUCTION

This paper is dedicated to the work and achievements of Watt W. Webb. Among his many important contributions to spectroscopy and microscopy are the development of Fluorescence Correlation Spectroscopy (FCS), together with Douglas Magde and Elliott Elson, (1–3), of two-photon excited fluorescence microscopy (4), together with Winfried Denk and James H. Strickler, and, the propagation of two-photon excitation FCS (5, 6), together with Petra Schwille and colleagues, which was first developed by Berland et al. (7). In all these methods, optical aberrations play an important role, because they lead to a change in the detection volume of a confocal microscope. In an FCS experiment, this results in a delayed decay of the correlation functions. Likewise, they will reduce the optical resolution of a two-photon microscope. The canonical way to describe optical aberrations caused by wavefront deformations in an optical system is to expand the wavefront deformations into an orthogonal set of functions over the unit disk, usually the Zernike polynomials. Frits Zernike, the inventor of the phase-contrast microscope (8) (which earned him the Nobel Prize in Physics in 1953), introduced these polynomials in 1934 (9). Their orthogonality makes them ideal for decomposing the wavefront error in optical systems into simpler, constituent aberrations. This decomposition enables an efficient understanding and correction of optical aberrations, which are critical for enhancing the quality of images produced by optical microscopy (10). Furthermore, Zernike polynomials play a crucial role in the development of adaptive optics systems for microscopy. These systems, which measure and actively compensate for wavefront errors, leverage devices like deformable mirrors or liquid crystal spatial light modulators. These devices are controlled based on the Zernike polynomial representation of aberrations, allowing for correction of distortions and the achievement of higher resolution and contrast images, especially in deep tissue imaging (11).

Although Zernike polynomials are well defined in many textbooks and research papers (12–14) and even on Wikipedia (15), the reasons behind their specific forms and the possibility of alternative function sets over the unit disk are less commonly known. This paper presents a derivation of Zernike polynomials based on variational calculus, providing insights into why they represent the simplest polynomial orthogonal function set over the unit disk. Along the way, we derive Bessel functions from a similar variational approach, noting that they are polynomial functions of infinite order.

In the second part of this paper, we will study the impact of different aberrations described by distinct Zernike functions on one-photon and two-photon excitation experiments. Although there have been extensive studies on the impact of various optical system imperfections on FCS experiments (see, e.g., Refs. (16, 17)), we are not aware of a comparative study between one-photon and two-photon FCS.

### ORTHOGONAL FUNCTIONS OVER THE UNIT DISK: BESSEL FUNCTIONS

An ideal microscope transforms an incoming spherical wavefront from a point source into an outgoing spherical wavefront that converges into a focus point in the image plane. However, real optical systems are never perfect and exhibit various deviations from this ideal behavior, collectively called aberrations. The most significant aberrations are due to an additional position-dependent *phase* of the wavefront. Although the field amplitude across the wavefront can also be affected by an optical system (through attenuation/apodization), this effect is usually less important because it does not alter the shape of the resulting point spread function as severely as phase variations do. Therefore, we will focus here solely on wavefront deformations. Conveniently, this phase is described by a system of orthonormal functions over the unit disk, where the unit disk maps the spherical wavefront of the non-aberrated system. Therefore, we aim to identify a set of orthogonal functions defined on the unit disk that is most suitable for describing aberrations.

Our goal is to find a linear differential equation whose eigenfunctions form a set of orthogonal functions over the unit disk. The canonical approach is to define a functional over the unit disk, where its minimization leads to the desired differential equation. Naturally, such a functional will be constructed using derivatives of the scalar function *f*, for which the differential equation is sought. The lowest possible derivative, which is also independent of the underlying coordinate system, is the gradient of *f*. However, the gradient returns a vector field, whereas the functional must be a scalar. Therefore, the simplest possible functional is constructed from the square (scalar product) of the gradient, i.e., it is given by the integral

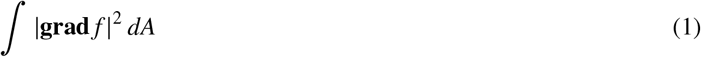

where *dA* represents the area element of the disk. In other words, we essentially target functions that show minimal deviation from being constant. Additionally, these functions should be normalized, which means the integral

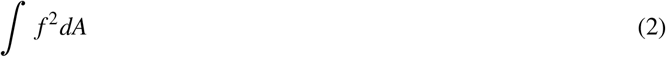

should equal one. By employing a Lagrange multiplier *λ*^2^, we can merge this normalization condition with the initial minimal bending energy requirement into seeking functions that minimize the functional

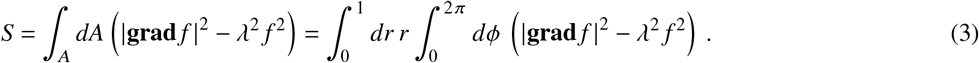

The simplest function meeting our criteria is a constant function. However, due to the orthogonality constraint, each subsequent function in our series must exhibit increasing complexity compared to all previously identified functions. This approach enables the construction of a sequence of functions where each new addition not only minimizes the mean-square gradient but also remains orthogonal to every preceding function, facilitating a systematic and unique expansion of the function space over the unit disk.

The variation of the above functional will lead to a linear partial differential equation for *f* in the variables *r* and *ϕ*, which can be analyzed and solved using the standard method of separation of variables. However, it is more convenient to perform the separation of variables before minimizing the functional (3) and to focus on the resulting differential equation for the radial part of *f*. Given that the functions *f* must exhibit 2*π* periodicity in the angular variable *ϕ*, we decompose *f* into *f* = *R* (*r*) cos *mϕ* or *f* = *R* (*r*) sin *mϕ*, where *R* (*r*) are real functions and *m* is an integer number. This decomposition leads, up to a constant pre-factor, to the following functional

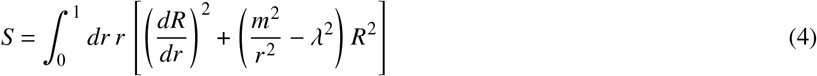

Variation of this functional yields

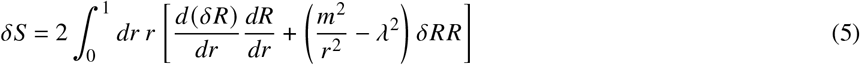

which, upon integration by parts, results in

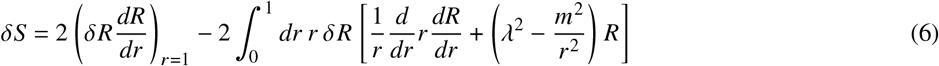

For the variation to disappear, the radial function *R*(*r*) must satisfy the differential equation

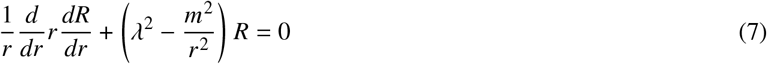

with the boundary condition *dR*/ *dr* = 0 at *r* = 1 ensuring that the partially integrated part in (6) disappears. Equation (7) is the Bessel differential equation, and its real and finite solutions *J*_*m*_ (*λr*) are the Bessel functions of the first kind *J*_*m*_.

Thus, the looked-for orthonormal function set on the unit disk is given by

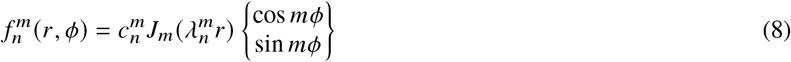

where the 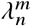 are the roots of the equation

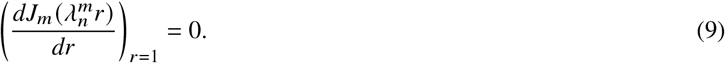

and 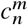 is a normalizing constant.

Let us check that two functions *J*_*m*_ (*λ*_*a*_ *r*) and *J*_*m*_ (*λ*_*b*_ *r*) with *λ*_*a*_ ≠ *λ*_*b*_ are indeed orthogonal to each other. We will use this method later also for Zernike polynomials. We first introduce the differential operator Ô defined by

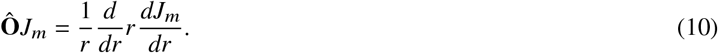

With this operator, the Bessel differential Equation (7) can be rewritten as

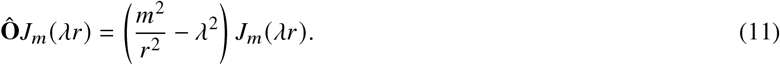

Application of this operator to function *J*_*m*_(*λ*_*a*_*r*) with left multiplication by *J*_*m*_(*λ*_*b*_*r*) yields

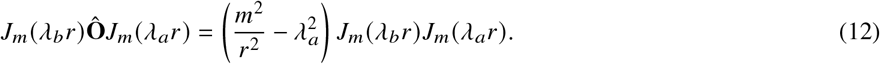

Subtraction of *J*_*m*_(*λ*_*a*_*r*) Ô *J*_*m*_(*λ*_*b*_*r*) and subsequent integration results in

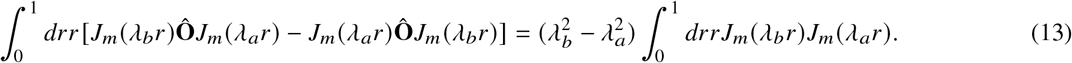

Integration by parts in the left hand side of the above equation leads to

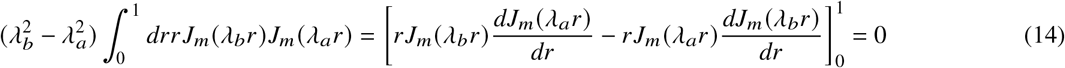

where we have used the fact that both *λ*_*a*_ and *λ*_*b*_ shall be roots of the equation [ *dJ*_*m*_ (*λr*) /*dr* _*r*=1_ = 0].

For our derivation of Zernike polynomials later, it is useful to recall the power series method for finding explicit solutions of differential equations such as the Bessel Equation (7). We expand the function *J*_*m*_(*r*) into a power series

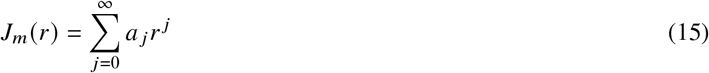

insert this into the differential Equation (7) with *λ* = 1, and compare the coefficients of equal power. This leads to the recursive relation

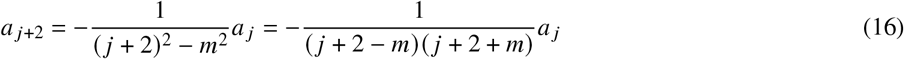

for the coefficients *a* _*j*_. This shows that the first non-vanishing coefficient must be *a*_*m*_ (to avoid infinite terms in the series of coefficients) and we find, after setting *a*_*m*_ = 1,

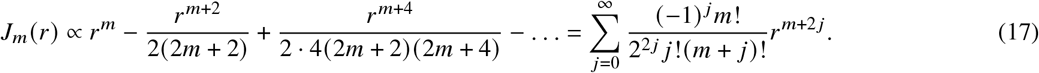

This leads, after canceling the common factor 2^*m*^*m*!, to the standard definition of the Bessel function *J*_*m*_(*r*)

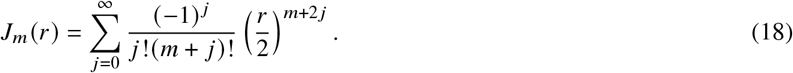

For finding its normalization, we consider the expression

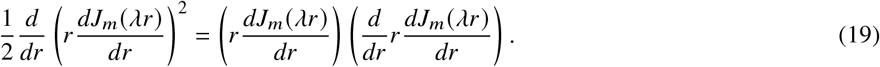

The second bracket on the r.h.s. can be transformed with Bessel’s differential Equation (7) to yield

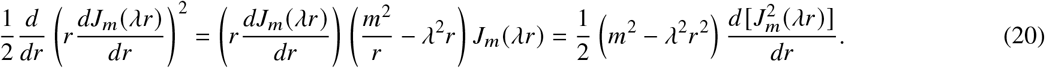

Thus, after integrating the l.h.s. and r.h.s of the last equation from *r* = 0 to *r* = 1, one obtains

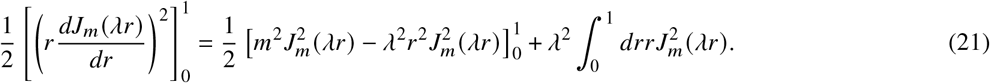

where we have employed once again a partial integration on the r.h.s. Consequently, when choosing for *λ* one of the roots 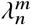 of [*dJ*_*m*_(*λr*)/*dr* = 0]_*r*=1_, and taking into account that *mJ*_*m*_(0) = 0 for all integer *m*, see (18), we find the normalization

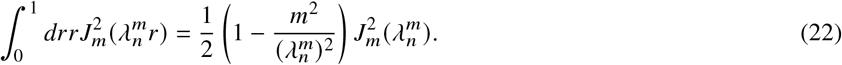

Thus, the normalizing constant 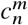 in (8) is

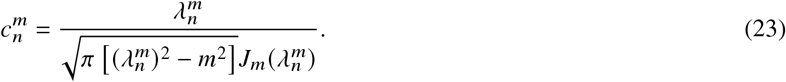

Figure 1 shows density plots of the first 12 modes 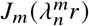 and 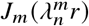 which are similar to the first 12 Zernike functions that describe the lowest order aberrations, see Figure 2 in the next section.

**Figure 1:**
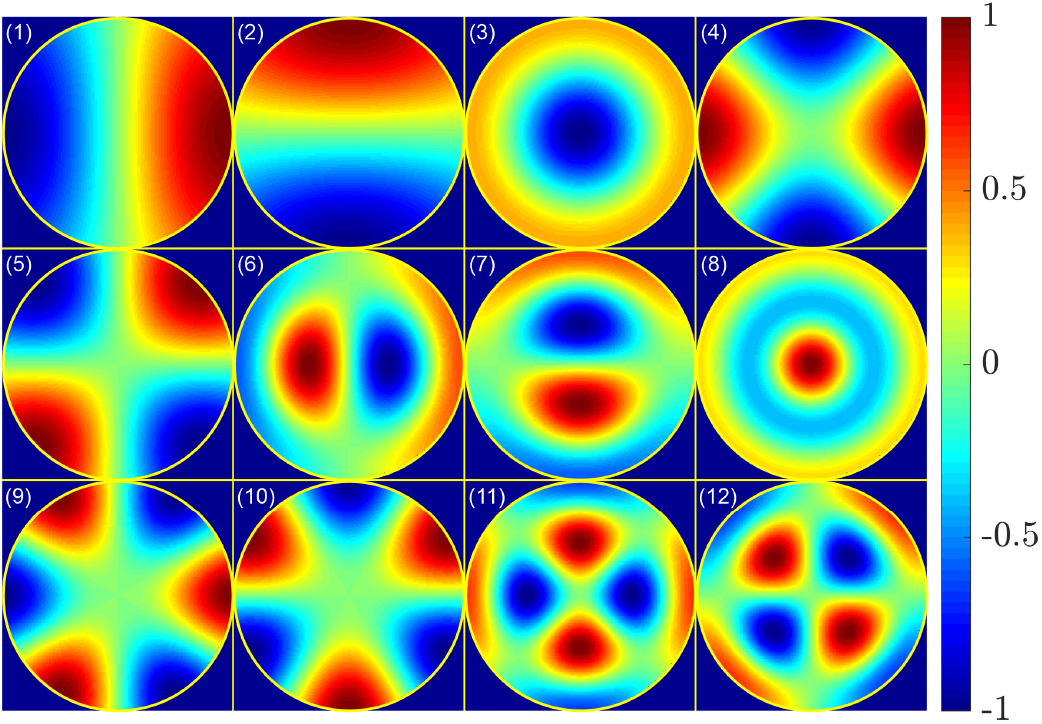
Density plots of the first low-order orthogonal functions 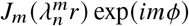;(1) 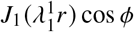; (2) 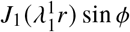; 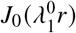; (4) 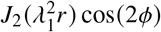; (5) 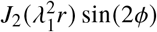 (6) 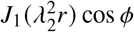; (7) 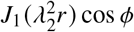; (8) 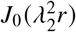; (9) 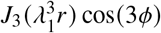; (10) 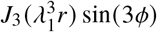; (11) 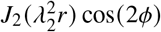; and (12) 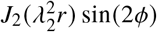. In the figures, all functions were normalized to their absolute maximum. The choice of the modes was done to make them most similar to the first canonical 12 Zernike aberrations, see Figure 2 and next section.

**Figure 2:**
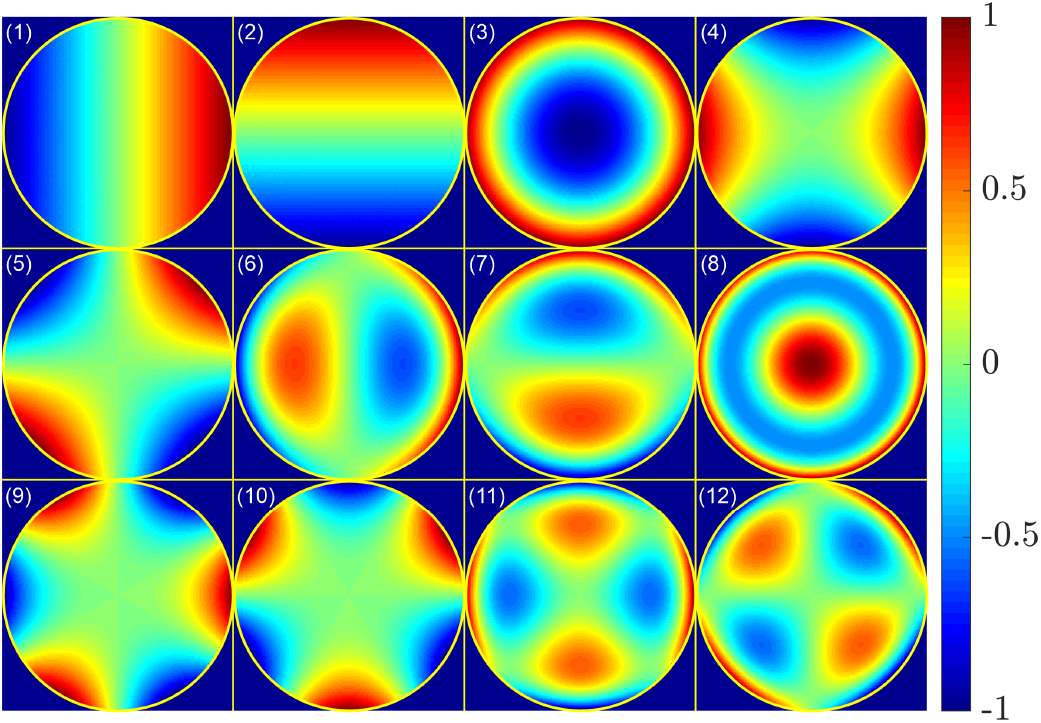
Density plots of the first twelve Zernike polynomials as presented in table 1: (1) horizontal or 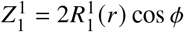; (2) vertical or tilt,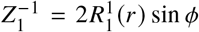; (3) defocus,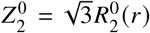; (4) vertical astigmatism,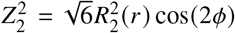;(5) oblique astigmatism, 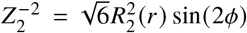; (6) horizontal coma, 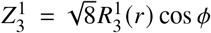; (7) vertical coma, 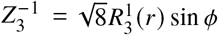; (8) primary spherical aberration, 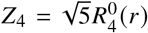; (9) oblique trefoil, 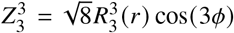; (10) vertical trefoil, 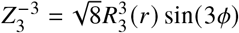; (11) vertical secondary astigmatism, 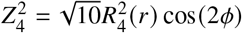); and (12) oblique secondary astigmatism, 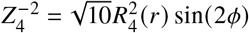.

### ZERNIKE POLYNOMIALS

As we have seen, Bessel functions can be used to construct an orthonormal set of functions over the unit disk, but their use has several drawbacks: Bessel functions are transcendental functions that have to be computed numerically, and even finding the roots of [*dJ*_*m*_ (*λr*) / *dr* = 0] _*r*=1_ is a nontrivial task. Although in the current era of powerful computers this is not a principal obstacle anymore, it was much more so in the times of Frits Zernike, who wanted to come up with a much simpler set of functions where the radial part can be represented by finite-order polynomials. From the power series method used to derive the series expansion of Bessel functions, it is evident that the Bessel differential equation cannot have finite-order polynomial solutions. This is because the differential operator reduces any power of *r* by two, while the term with *λ*^2^ does not. As a result, the recurrence relation for the series expansion coefficients can never terminate at a finite power of *r*. To rectify this situation, one may introduce into the functional *S* another differential operator which does *not* lower the order of a polynomial but still is *rotation-invariant*. Such an operator is div(**r** *f*), so that we may use the modified functional

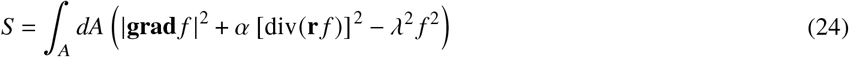

with some constant *α* as the starting point. As can be checked later, the alternative choice of **r · grad** *f* instead of div (**r** *f*) gives the same final result for the differential equation. Although it is not guaranteed that the modified functional (24) will lead to a differential equation of *f*, we again use the Ansatz *f* = *R*(*r*) cos *mϕ* or *f* = *R*(*r*) sin *mϕ* which now leads to

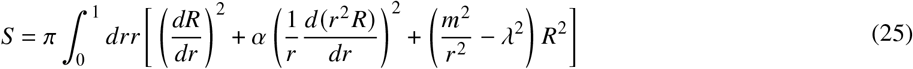

**Table 1:** The first 12 Zernike functions.

| # | $n$ | $m$ | function name | $R_n^m(r) \begin{cases} \cos m\phi \\ \sin m\phi \end{cases}$ | name |
| --- | --- | --- | --- | --- | --- |
| 1 | 1 | 1 | $Z_1^1$ | $2r \cos \phi$ | horizontal tilt |
| 2 | 1 | -1 | $Z_1^{-1}$ | $2r \sin \phi$ | vertical tilt |
| 3 | 2 | 0 | $Z_2^0$ | $\sqrt{3}(2r^2 - 1)$ | defocus |
| 4 | 2 | 2 | $Z_2^2$ | $\sqrt{6}r^2 \cos 2\phi$ | vertical astigmatism |
| 5 | 2 | -2 | $Z_2^{-2}$ | $\sqrt{6}r^2 \sin 2\phi$ | oblique astigmatism |
| 6 | 3 | 1 | $Z_3^1$ | $\sqrt{8}(3r^2 - 2)r \cos \phi$ | horizontal coma |
| 7 | 3 | -1 | $Z_3^{-1}$ | $\sqrt{8}(3r^2 - 2)r \sin \phi$ | vertical coma |
| 8 | 3 | 3 | $Z_3^3$ | $\sqrt{8}r^3 \cos 3\phi$ | oblique trefoil |
| 9 | 3 | -3 | $Z_3^{-3}$ | $\sqrt{8}r^3 \sin 3\phi$ | vertical trefoil |
| 10 | 4 | 0 | $Z_4^0$ | $\sqrt{5}(6r^4 - 6r^2 + 1)$ | primary spherical |
| 11 | 4 | 2 | $Z_4^2$ | $\sqrt{10}(4r^2 - 3)r^2 \cos 2\phi$ | vertical secondary astigmatism |
| 12 | 4 | -2 | $Z_4^{-2}$ | $\sqrt{10}(4r^2 - 3)r^2 \sin 2\phi$ | oblique secondary astigmatism |

Variation and partial integration of this functional leads to

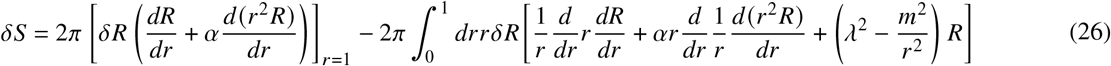

To make the partially integrated term at *r* = 1 independent on the derivative of *R*, one sets *α* =−1, so that the above equation simplifies to

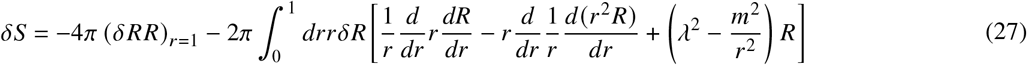

which leads to the ordinary differential equation

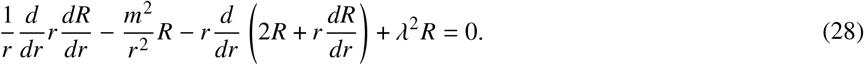

for *R*(*r*). When we use a finite-order polynomial Ansatz for *R*(*r*) with highest order *n*, we see that this can indeed work if *λ*^2^ = *n*(*n* + 2), and if the coefficients *a* _*j*_ for *j* < *n* obey the recursive relation

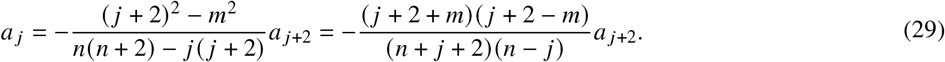

Thus, we find for the radial polynomials 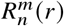 the explicit expressions

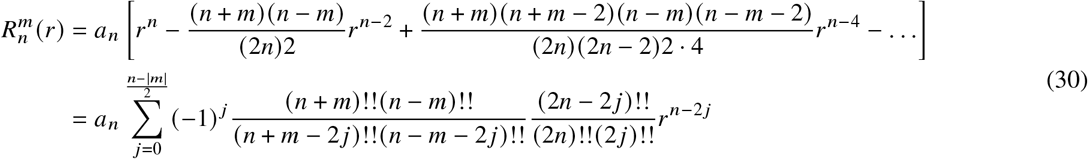

where *x*!! denote the double factorial *x*!! = *x* (*x* − 2)(*x* − 4) · · ·. The index *m* in 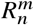 describes the periodicity of the final function 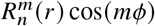 or 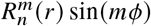 in the angular variable *ϕ*, but *n* is now the polynomial order of the radial function. Importantly, we see that *n* − *m* must be even so that no negative powers of *r* occur in the above expansion. When taking into account that for even integer numbers *k* one has *k*!! = 2_*k*/2_ (*k*/2)!, one finds

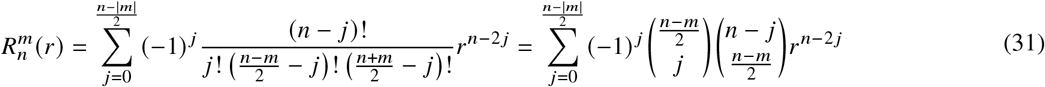

where we dropped the highest coefficient *a*_*n*_ and all factors that do not depend on the summation variable *j*.

Next, we want to calculate the value of 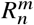 at *r* = 1. For doing this, consider the *q*^th^ power (*x*− *y*)^*q*^ of the difference between two auxiliary variables *x* and *y* and expand it into a binomial series

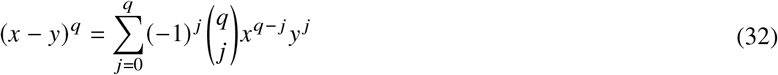

When multiplying both sides of this equation with *x*^*n*−*q*^, differentiating *q* time with respect to *x*, and dividing by *q*!, we find

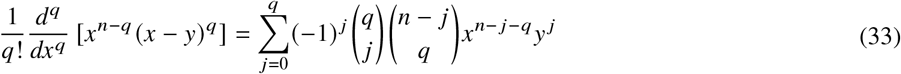

However, when using the binomial formula for expanding the *q*^th^ derivative on the l.h.s., we find

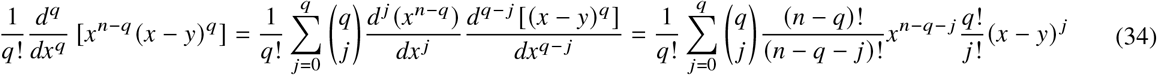

When setting in both previous equations *x* = *y* = 1, this yields

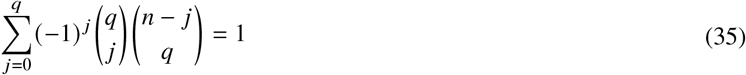

which shows that 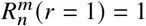. Thus, all radial functions 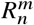 are constant and equal to one along the circumference of the unit disk (*r* = 1), so that we have indeed *δR*| _*r*=1_ = 0 in the variation (27).

To establish the orthogonality of the Zernike polynomials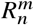, we start by acknowledging that the orthogonality of 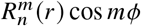 and 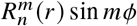 for differing *m* values is secured by the intrinsic orthogonality of the trigonometric functions cos *mϕ* and sin *mϕ*. Our focus now shifts to demonstrating the orthogonality of 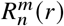 across different *n* values while maintaining *m* the same. For doing that we abbreviate the differential operator in (28) by Ô as in the previous section, i.e.

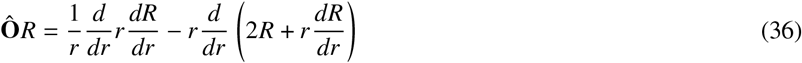

so that, when using the differential Equation (28), we have

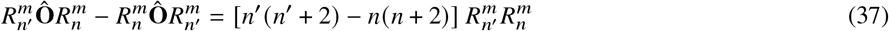

On the other hand, we obtain after partial integration,

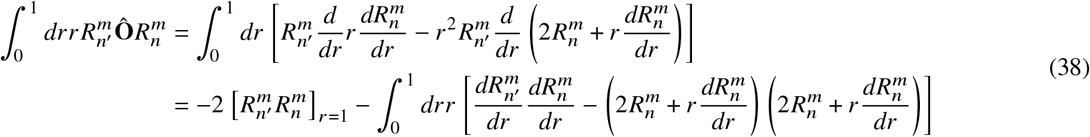

so that

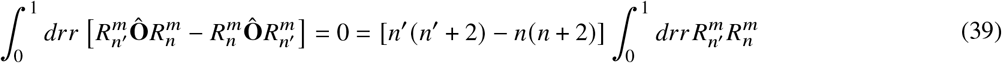

which proves the orthogonality of the 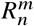 for different values of *n* but same *m*.

Finally, we want to find the normalization of the Zernike polynomials, i.e. we want to calculate the integral 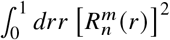.

Here, we can use the fact that the 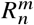 are *n*^th^ order polynomials that are orthogonal to each other. Thus, the part of 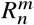 that contains all the powers of *r* smaller than *n* can be expressed by an expansion into polynomials 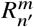 with *n*^′^ < *n*. Due to the orthogonality of the Zernike polynomials and using the explicit expression (31), we thus find

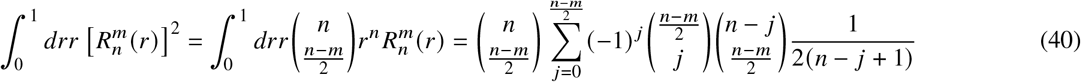

To simplify the r.h.s. of this equation, we consider the integral

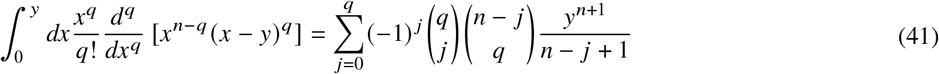

where the r.h.s. was found by expanding (*x* − *y*)^*q*^ into the sum 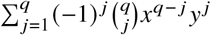 and then differentiating and integrating each term of this sum separately. However, we can also perform *q* times a partial integration on the l.h.s., which leads to the alternative result

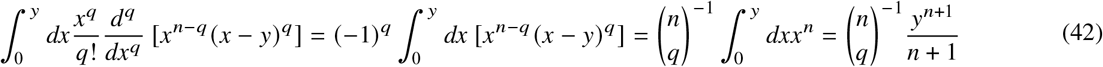

where we have, for coming from the second to the third term, applied *q* times a partial integration to the powers of *x*. By equating the r.h.s. of (41) and (42), we have

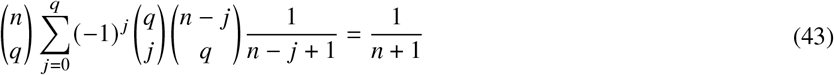

and then comparing the l.h.s. of this expression with (40), we finally find

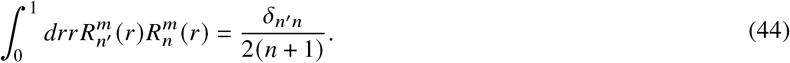

The complete formulation of Zernike functions is concisely captured by the equation:

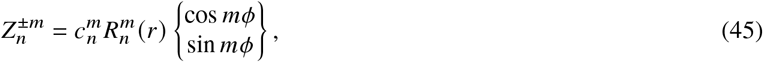

where in the superscript ±*m* distinguishes between the cosine function (for a positive *m*) and the sine function (for a negative *m*). The normalization coefficient 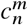 is selected to ensure that the integral over the unit disk is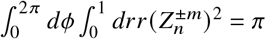. Using the normalization detailed in (44), this leads to the explicit definition of 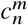 as:

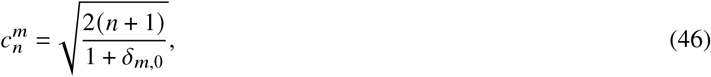

which standardizes the Zernike polynomials across the unit disk, facilitating their application in optical systems for aberration correction and wavefront analysis.

## ABERRATIONS, FOCUS SHAPE, AND FLUORESCENCE CORRELATION SPECTROSCOPY

In this section, we consider the impact of aberrations as described by Zernike functions on the focusing of light through an objective, and the consequences for fluorescence correlation spectroscopy using a confocal microscope. In the first subsection, we will consider focusing of a plane wave, in the second subsection, the detection efficiency distribution of confocal imaging, and in the third subsection, how aberrations affect the outcome of an FCS measurement.

### Focusing

When focusing a linearly polarized plane wave (polarization direction along the *x*-direction in Figure 3) with field amplitude *E*_0_ through an objective into a sample with refractive index *n*_*r*_, the electric field distribution close to the focus position **r** = 0 = {0, 0, 0} is given by (18, 19)

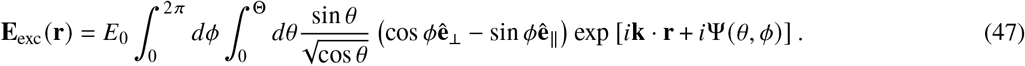

which represents a superposition of plane waves with wave vectors **k**. The meaning of the involved vectors **k, ê**_⊥_ and **ê**_∥_ is visually explained in Figure 3(a). The factor 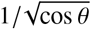 ensures conservation of energy flux along the optical axis while the light is focused through the objective. Abbe’s sine condition relates the distance *r* from a point on the incident planar wavefront to the angle *θ* at a corresponding point on the focused spherical wavefront through the relationship *r* = *f* sin *θ*. Here, *f* denotes the focal length of the lens focusing the wavefront. The upper integration limit Θ is the half-angle of light collection of the focusing objective, which is related to its numerical aperture via NA = *n* sin Θ.

**Figure 3:**
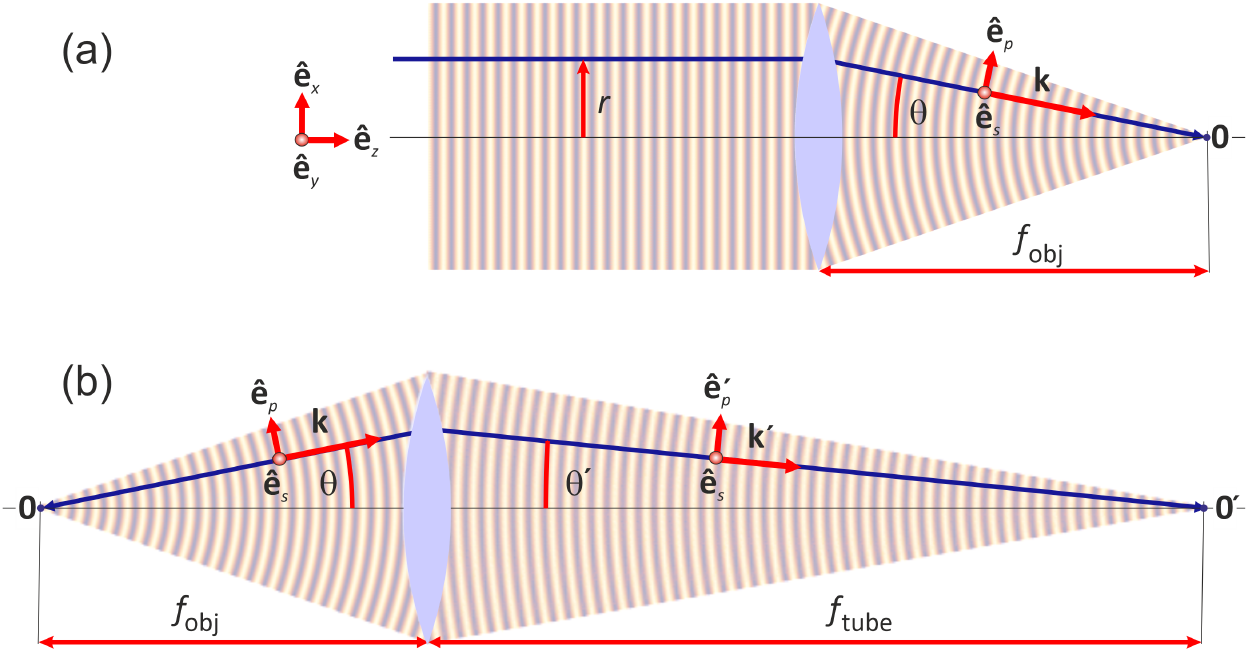
Excitation and detection in a FCS experiment. (a) Focusing of a linearly polarized plane wave (polarization along vector **ê**_*x*_) through an objective lens with focal length *f*_obj_. The electric field close to the focal point 0 can be represented as a superposition of plane waves traveling along all possible wave vectors **k** = *nk*_0_ sin *θ* cos *ϕ*, sin *θ* sin *ϕ*, cos *θ*, where *n* is the refractive index of the medium where the light is focused, and *k*_0_ = 2*π λ* is the length of the vacuum wave vector at wavelength *λ*. The polarization of each plane wave can be divided into two components along the unit vectors **ê**_*p*_ = cos *θ* cos *ϕ*, cos *θ* sin *ϕ*, sin *θ* and **ê**_*s*_ = sin *ϕ*, cos *ϕ*, 0. (b) Wave optics of imaging a point source (electric dipole emitter) in sample space (left hand side) into the plane of the confocal pinhole (right hand side). Each plane wave component with wave vector **k** = *nk*_0_ {sin *θ* cos *ϕ*, sin *θ* sin *ϕ*, cos *θ*} in sample space is converted into a plane wave with wave vector **k**^′^ = *k*_0_ {− sin *θ*^′^ cos *ϕ*, − sin *θ*^′^ sin *ϕ*, cos *θ*^′^} in image space. The relation between *θ* and *θ*^′^ is given by Abbe’s sine condition, *n* sin *θ* = *M* sin *θ*^′^, where *M* = *f*_tube_/ *f*_obj_ is the imaging magnification. While the unit vector for the *s*-wave **ê**_*s*_ = {− sin *ϕ*, cos *ϕ*, 0} is the same in the sample and the image space, the unit vector for the *p*-wave in sample space is **ê**_*p*_ = {cos *θ* cos *ϕ*, cos *θ* sin *ϕ*, − sin *θ*} and in image space is 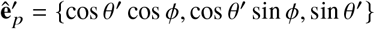.

The function Ψ (*r, ϕ*)describes wave front deformations of the incoming plane wave and/or ones induced by an imperfect focusing optics. These wavefront deformations result in aberrations of focus shape and alter the light intensity distribution across the focus. The function Ψ(*r, ϕ*) is efficiently represented by a series expansion in terms of Zernike functions:

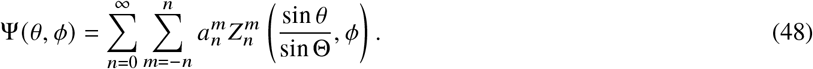

The coefficients 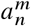 are the expansion coefficients that quantify the contribution of each Zernike function to the overall wavefront distortion. We now have everything in place to delve into the effects of various aberrations on the shape and size of the laser light focus. The lowest non-trivial Zernike functions, specifically 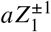 with an amplitude factor *a*, represent linear gradients along the *x*- or *y*-axis. Incorporating these linear expressions into the exponent on the r.h.s. of (47) reveals that they induce a lateral displacement of the focal position by 2*a*/(*k* sin Θ), without modifying its shape. In asimilar vein, the aberration denoted by 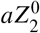 leads to a first-order shift of the focus along the optical axis, quantified by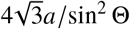. Hence, these initial Zernike functions effectively capture simple translational shifts of the focus, illustrating their superior utility in optical system descriptions as opposed to the orthogonal function system that involves Bessel functions. The latter, depicted in Figure 1, which parallel the Zernike expansion modes presented in Figure 2, convey a considerably more intricate physical interpretation. For instance, the primary three Bessel modes, corresponding to straightforward focus shifts in the Zernike framework, also impact the focus’ shape, as they cannot be succinctly represented by simple first or second-order polynomials in the radial coordinate. In Figure 4, we examine the influence of the lowest significant Zernike modes on the shape of the focus. Specifically, we investigate modes that characterize first and second order astigmatism, coma, primary spherical aberration, and trefoil. In each instance, we have set the amplitude of the corresponding Zernike functions to unity. Astigmatism is typically caused by non-planar surfaces of reflective elements such as beam splitters, dichroic mirrors, or plane mirrors. Coma is most often introduced by later misalignment of lens elements with respect to the optical axis. Primary spherical aberration is primarily caused by the classical spherical aberration of spherical (non-parabolic) lenses, although high-quality optical systems are quite effective at compensating for these effects. Finally, trefoil aberrations can be induced by diffraction from obstructions within the optical pathway, such as the three-armed holders used for optical elements in mirror telescopes.

**Figure 4:**
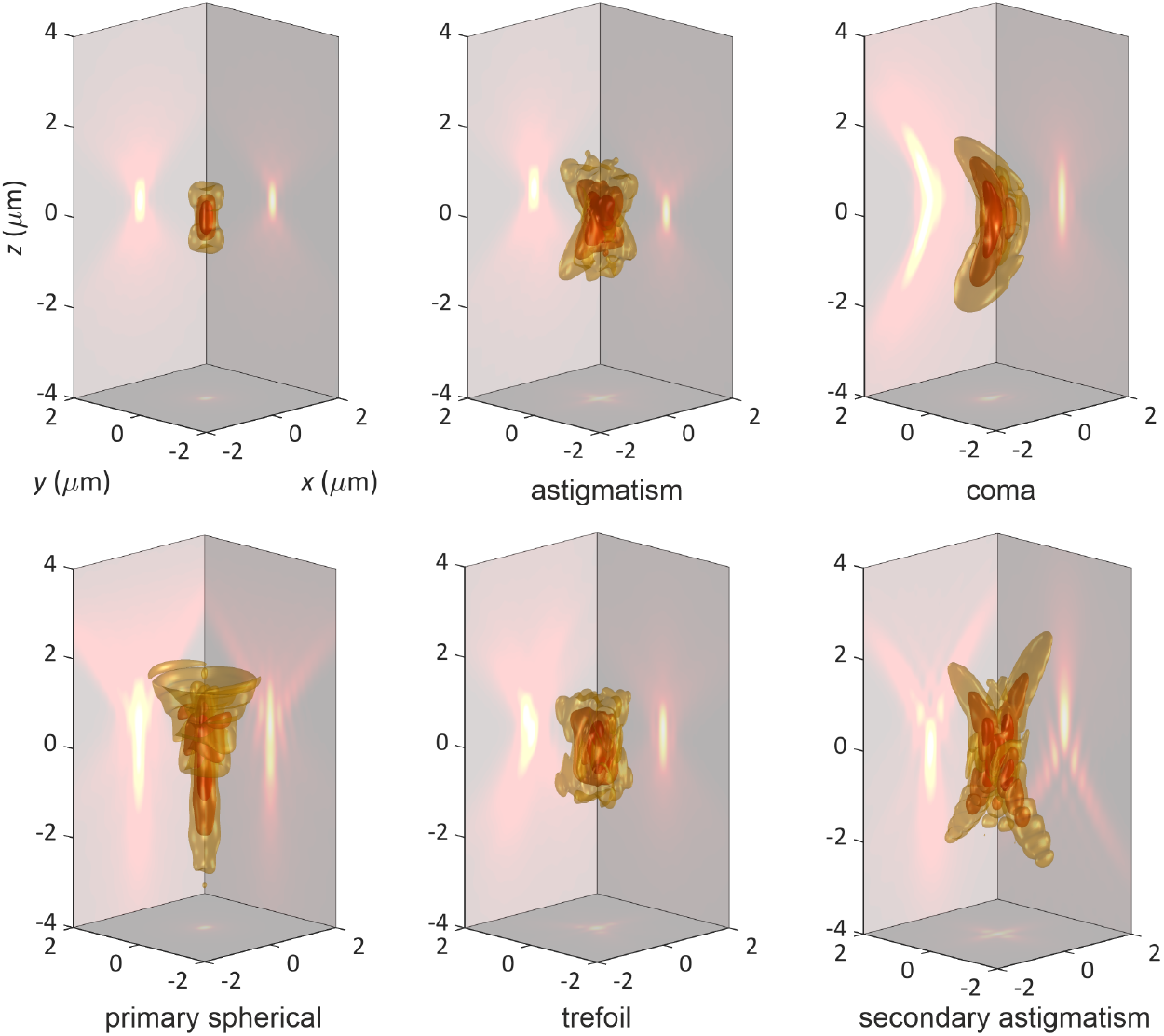
Iso-surface plots of *U* (**r**) _exc_ = |**E** (**r**)| ^2^ (shown are the surfaces where the intensity *U*_exc_ has fallen to 1/ *e*, 1 /*e*^2^, and 1 /*e*^3^ of its maximum value in the center) and maximum intensity projections of *U* for different types of aberrations. Calculations were done for focusing an *x*-polarized plane wave with 532 nm wavelength through a water immersion objective with a numerical aperture of NA = 1.2. The top left panel shows the ideal aberration-free case. All other panels show the effect of different aberrations as indicated below each panel, and all axes values are in μm as shown in the first panel.

### Confocal detection

In an epi-fluorescence confocal microscope, as typically used in conventional FCS, the fluorescence excited within the focused laser light is collected by the same objective and then focused through a confocal aperture before being detected by a point light detector. This is done to ensure efficient rejection of out-of-focus light and thus to generate a detection volume with its lateral site determined by the diameter of the focused laser light, and its axial size by the confocal detection.

Basically all common fluorescent dyes, proteins or quantum dots behave as ideal electric dipole emitters. Thus, for modeling the detection efficiency distribution of a confocal microscope, we consider the imaging of an electric dipole emitter in sample space through the objective into the plane of the confocal pinhole. The electric and magnetic field distributions at lateral position *ρ* in that plane generated by an electric dipole emitter (fluorescent molecule) with dipole moment amplitude **p** at position **r** in sample space are given by (see for example (20))

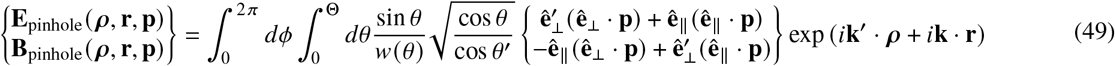

where the meaning of all vectors **k, k′**, **ê**_⊥_, 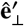 and **ê**_∥_ is visually explained in Figure 3(b). The factor 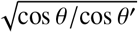 in (49) takes again care of conservation of the energy flux along the optical axis, and *w*(*θ*) = *nk*_0_ cos *θ* is the axial component of the wave vector **k** in sample space. Next, the detection efficiency through the confocal pinhole for such an emitter is given by the integral of the time-averaged axial component of the Poynting vector, **ê**_*z*_ · **P** = (*c*/8*π*)**ê**_*z*_ · (**E**_pinhole_ × **B**_pinhole_), over the pinhole area,

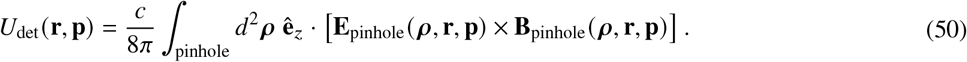

Finally, for an isotropic distribution of molecule orientations, the detection efficiency is obtained by averaging the above expression over all orientations of the electric dipole vector 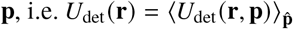. For the sake of simplicity, when modeling an FCS measurement in the next subsection we assume that aberrations are only present in the focusing of the fluorescence excitation light, but not in the imaging from sample to image space. However, this is no general restriction–imaging aberrations could be treated by including a similar phase function Φ (*θ, ϕ*) into the exponent in (49) as was done in (47) when considering aberrations in excitation light focusing.

### Fluorescence correlation spectroscopy

FCS is a single-molecule sensitive technique that measures and evaluates the intensity fluctuations of the fluorescence that is excited within the minuscule detection volume of a confocal microscope (1–3, 21). The autocorrelation function calculated from these intensity fluctuations reveals insights into any physical processes that influence the fluorescence intensity fluctuations within this volume. A primary application of FCS is to study the translational diffusion of fluorescent molecules through the detection volume, providing data on the diffusion coefficients of these molecules. However, the diffusion-related decay time of the autocorrelation function is notably sensitive to the size and shape of this detection volume (16, 17). Consequently, various FCS adaptations have been developed to offer robustness against potential optical aberrations that might occur in a confocal microscope setup. Among these methods, notable examples include dual-focus FCS (2fFCS) (22, 23), scanning FCS (24, 25), and z-scan FCS (26, 27). Two-photon excitation FCS (5–7) also promises to minimize impact from optical aberrations, thanks to the quadratic intensity dependence of fluorescence excitation. This dependency selectively emphasizes the contributions from the focus’s high-intensity central region over its lower-intensity surrounding areas. In this study, we utilize Zernike functions to model optical aberrations incurred while focusing a laser beam through an objective and assess how these aberrations influence one-photon and two-photon excitation FCS experiments.

The excitation intensity distributions for one- and two-photon excitation are determined by the second and fourth absolute power of the electric field distribution described by (47). In case of one-photon excitation FCS, the excitation intensity distribution has to be multiplied with the detection efficiency distribution *U*_det_ (**r**)of (50), i.e. the three-dimensional function that describes how well an emitting molecule is detected through the pinhole of the confocal microscope, to obtain the total molecule excitation and detection probability distribution. In case of two-photon excitation FCS, which does not require confocal detection (due to its intrinsic optical sectioning capability), this excitation and detection probability distribution is given by the square of the excitation intensity distribution directly.

With the excitation and detection probability distributions determined, the diffusion-related component of the autocorrelation function, *g* (τ), is defined as the average of the product of three probabilities: the probability of exciting and detecting a fluorescence photon emitted by a molecule initially at position **r**_1_, the probability that this molecule diffuses to position **r**_2_ within the lag time τ, and the probability of subsequently exciting and detecting another photon from the molecule at its new position. This concept is mathematically represented by a six-dimensional integral

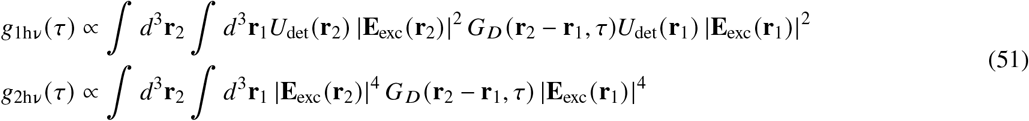

where *G*_*D*_ (**r**_2_ − **r**_1_, τ) is Green’s function for the free diffusion equation,

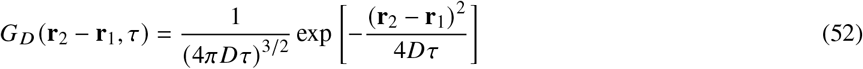

with *D* being the diffusion coefficient of the diffusing fluorescent molecules.

For the numerical calculation of this autocorrelation function, we first determined the electric field distribution **E**_exc_ (**r)** and the detection efficiency distribution *U*_det_ (**r**) across a grid of finite elements, each measuring 10× 10× 20 nm^3^, within a volume of 4× 4× 8 μm^3^. The calculations were performed with the following parameter values: one-photon excitation wave length of 532 nm, two-photon excitation wavelength of 1064 nm, water immersion objective with an NA of 1.2, aqueous sample medium with refractive index of 1.33, magnification of imaging 60 times, and pinhole diameter of 86 μm (corresponding to the position of the second Airy ring and assuring that 97% of light from an emitter at the objective’s focus that hits the plane of the pinhole is transmitted through the pinhole). Next, we computed the convolution ∫ *d*^3^r_1_*G*_*D*_(r_2_ ™ r_1_, *τ*)*U*^det^ (r_1_) |E_exc_ (r_1_) |^2^ for the one-photon excitation FCS and ∫ *d*^3^r_1_*G*_*D*_(r_2_ ™ r_1_, *τ*) |E_exc_ (r_1_) |^24^ for the two-photon excitation FCS in (51), respectively, employing Fast Fourier Transforms (FFTs) (28). This approach leverages the principle that the convolution of two functions in real space is proportional to the inverse Fourier transform of the product of their individual Fourier transforms. Finally, we achieved the second integration in (51) by aggregating the products of the convolution and *U* across all finite elements. Finally, we multiplied the results of these convolutions with either *U*_det_ (**r**_2_) |**E**_exc_(**r**_2_) | for the one-photon excitation case or with |**E**_exc_(**r**_2_) | for the two-photon excitation case and integrated the result over the coordinate **r**_2_. This yields the value of the corresponding autocorrelation function at lag time τ.

Figure 5 shows both the explicit autocorrelation curves for the ideal and different aberrated detection volumes and the resulting diffusion times defined by the lag time when the autocorrelation function has fallen to half of its maximum value. One observes that different types of aberrations change the shape of the FCS autocorrelation curve in different ways. For example, secondary astigmatism changes the curve shape in such a way that it is sufficiently distinct from that of an aberration-free experiment, which would help to discern it as coming from a measurement with substantial aberration. However, trefoil mostly shifts the correlation curve to longer times without much changing its shape, so that such a measurement could be easily confused with an aberration-free measurement of a slower diffusing sample. This is a principal problem of conventional FCS when using it for precise measurements of diffusion coefficients: It is very sensitive to optical aberrations, and its is hard to detect this by just inspecting FCS curve shape and resulting fit quality (16, 17). Comparison of the diffusion times with that of an ideal aberration-free measurement shows that, except for astigmatism, two-photon excitation FCS is slightly more robust against optical aberrations than one-photon excitation FCS. However, the difference between the performance of one- and two-photon excitation FCS is not as striking as one may hope for, emphasizing again the importance of more accurate FCS variants mentioned in the beginning of this section (22–27) when one wants to measure diffusion coefficients with high accuracy.

**Figure 5:**
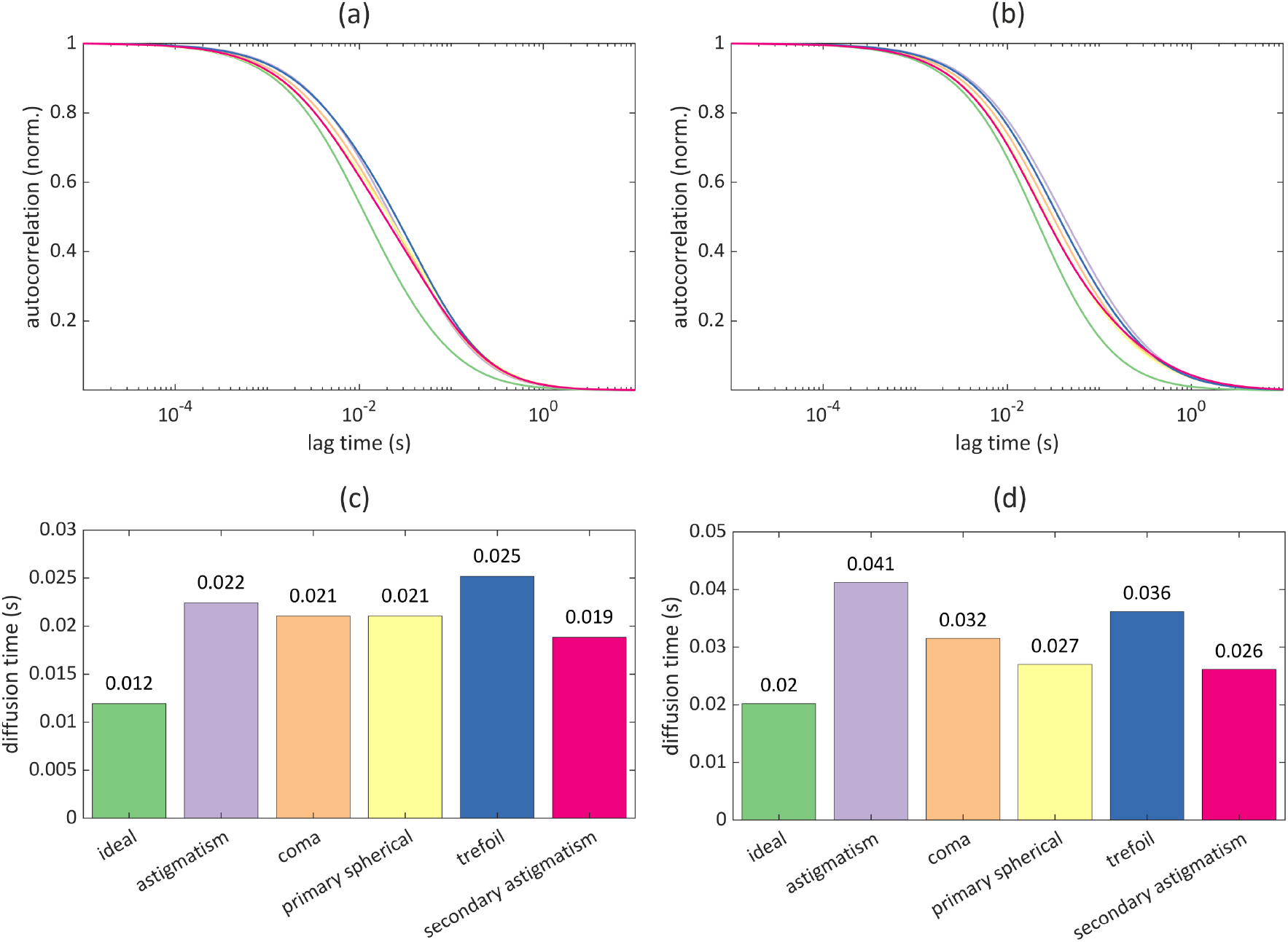
Fluorescence Correlation Spectroscopy (FCS) with aberrations. (a) Modeling of one-photon excitation FCS curves for pure diffusion through the ideal and the five aberrated detection volumes corresponding to the intensity distribution functions shown in Figure 4. The FCS curve for the ideal aberration-free detection volume is the leftmost curve. (b) Same as panel (a), but for two-photon excitation FCS. (c) Diffusion times (lag time when the autocorrelation has fallen off to half its maximum value) corresponding to the autocorrelation curves shown in panel (a). The colors of the bars correspond to the lines of the FCS curve plots, values above each bar denote the diffusion time in seconds. (d) Same as panel (c), but for two-photon excitation FCS.

Another crucial parameter in an FCS measurement is the maximum brightness of a molecule, defined as the brightness measured for a molecule positioned at the center of the focus and on the optical axis. Figure 6(a,b) shows this value compared to that observed for an ideal system without aberrations. It is evident that aberrations have a much more dramatic impact on maximum brightness than on diffusion time, and this effect is significantly worse for two-photon excitation experiments compared to one-photon excitation experiments. This observation correlates well with the impact of aberrations on the detection volume, which will be discussed next.

**Figure 6:**
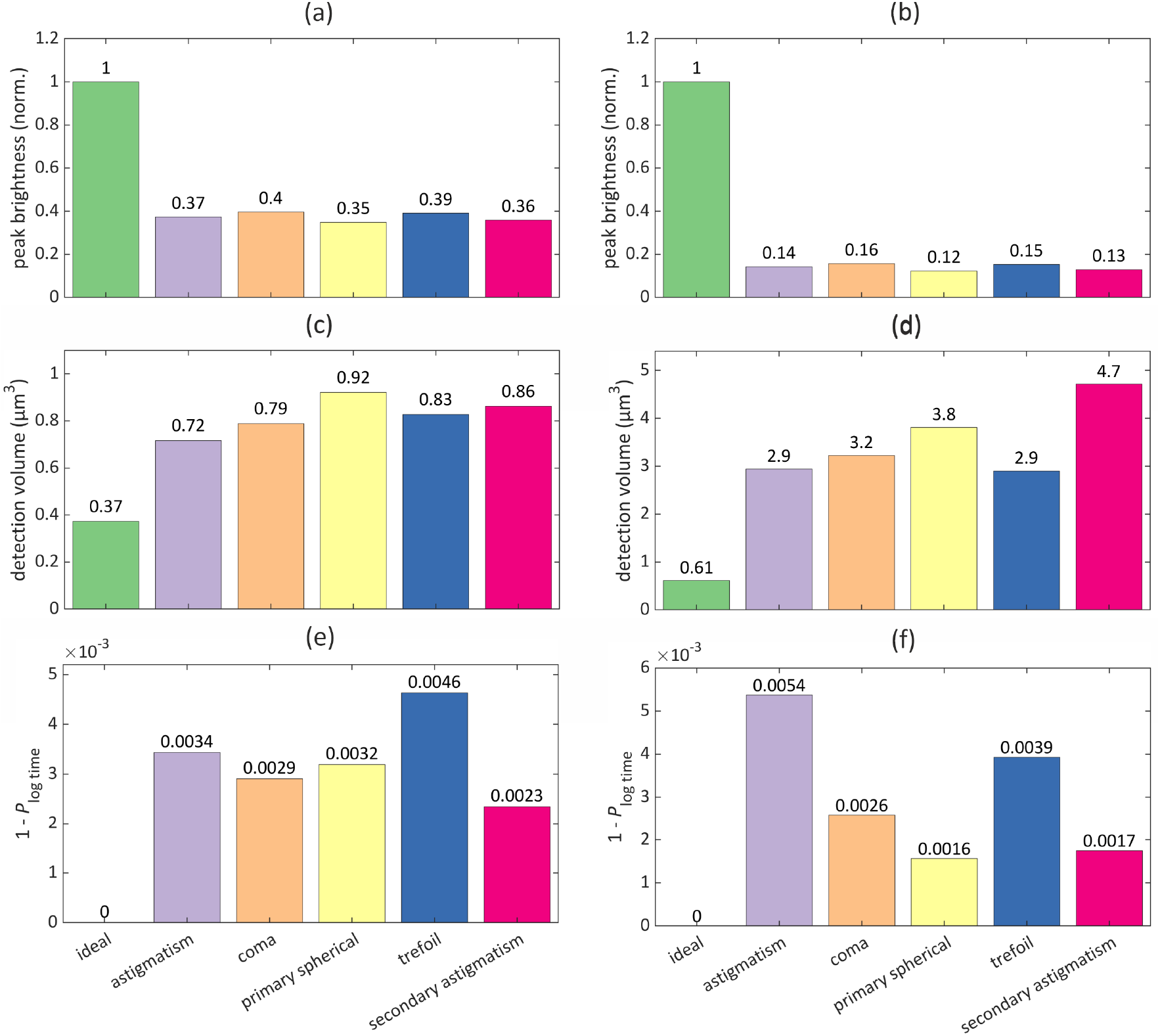
Brightness, detection volume, and curve similarity. Left panels show the result for 1-photon excitation FCS, right panels for two-photon excitation FCS. All model parameters are the same as used for Figure 5. (a) and (b) show the brightness that would be measured from a molecule in the focal plane and on the optical axis, as compared to a molecule a the same position in a measurement without any aberrations. (c) and (d) Effective detection volumes, see eq. (53). (e) and (f) Reduced logarithmic-time Pearson coefficients, see eq. (54), which are a measure of the similarity between autocorrelation curves measured with and without aberrations.

The detection volume *V*_det_ is a crucial characteristic of an FCS experiment when estimating the concentration of diffusing molecules. Its product with the concentration is inversely proportional to the relative amplitude of an FCS curve, given by lim 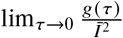 where 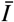 is the average measured fluorescence intensity. Thus, by measuring the relative FCS amplitude and knowing the detection volume, one can calculate the absolute concentration of fluorescent molecules in a sample. Mathematically, *V*_det_ is defined by

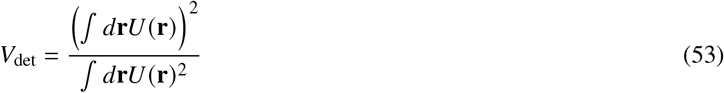

where we have *U*(**r**) = *U*_det_ (**r**) |**E**_exc_(**r**) |^2^ for one-photon excitation FCS, and *U*(**r**) = |**E**_exc_(**r**) |^4^ for two-photon excitation FCS. The computational results for the detection volume are shown in Figure 6(c,d). As illustrated, the impact of aberrations on *V*_det_ is similarly severe to that on brightness, and it is surprisingly more pronounced for two-photon excitation than for one-photon excitation. This behavior was previously reported in Ref. (17), which also noted a much stronger impact of aberrations on FCS-based estimates of concentration compared to diffusion coefficients (for one-photon excitation FCS).

Finally, to characterize the similarity (or lack thereof) between autocorrelation curves measured with and without aberrations, we use the reduced logarithmic-time Pearson coefficient 1 − *p*_log time_. Here, *p*_log time_ is defined as:

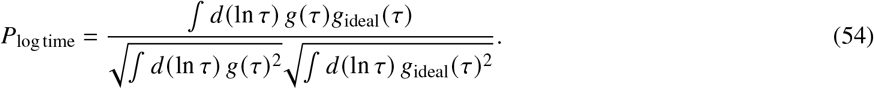

Calculating this coefficient with logarithmic time accounts for the rapid decay of an autocorrelation curve over time, as seen in the logarithmic plots in Figure 5(a,b), so most of its information is contained at short timescales. The reduced Pearson coefficient 1− *p*_log time_ will be exactly zero when comparing identical curves, and its absolute value increases with the difference between two curves. The computational results for this coefficient are shown in Figure 6(e,f). As illustrated, the difference between aberrated and ideal autocorrelation curves, as measured by 1− *p*_log time_, is similar across all considered aberration types for one-photon excitation, with trefoil aberration showing the most significant impact. However, for two-photon excitation FCS, astigmatism has by far the largest impact, causing the strongest deviation of the autocorrelation curve from its ideal, aberration-free case. Please note the qualitatively similar trends in the impacts of aberrations on diffusion time and on the similarity between autocorrelation curves.

## CONCLUSION

In this paper, we have presented a derivation of Zernike polynomials using a variational calculus approach and have derived the fundamental properties regarding orthogonality and normalization. On the way, we have elucidated the connection between Bessel functions and Zernike polynomials. Finally, we have applied Zernike functions for modeling the focusing of light through a high-aperture objective and then used this result for modeling the impact of different aberrations on one and two-photon excitation FCS measurements.

## CODE AVAILABILITY

The MATLAB code for performing all model calculations described in this paper is freely available from the authors upon reasonable request.

## AUTHOR CONTRIBUTIONS

Jörg Enderlein developed the general idea of the manuscript, wrote its core part, performed all numerical calculations and prepared all figures. Ivan Gligonov helped with editing the manuscript and checked all mathematical derivations and equations.

## DECLARATION OF INTERESTS

The authors declare no competing interests.

## ACKNOWLEDGMENTS

Jörg Enderlein acknowledges the financial support of the European Research Council (ERC) for financial support through project “smMIET” (grant agreement no. 884488) under the European Union’s Horizon 2020 research and innovation program, and financial support from the Deutsche Forschungsgemeinschaft through the German Excellence Strategy EXC 2067/1-390729940. Ivan Gligonov acknowledges funding by the International Max Planck Research School for Physics of Biological and Complex Systems and by the European Union via the HORIZON–MSCA–2022–DN “Improving BiomEdical diagnosis through LIGHT-based technologies and machine learning – BE-LIGHT” (GA n° 101119924 – BE-LIGHT).

## Notes

### Competing Interest Statement

The authors have declared no competing interest.

### Summary of Updates

Extended discussion of fluorescence correlation spectroscopy, additional figures.

## REFERENCES

1. Magde, D., E. Elson, and W. W. Webb, 1972. Thermodynamic fluctuations in a reacting system—measurement by fluorescence correlation spectroscopy. Physical Review Letters 29:705.

2. Magde, D., E. L. Elson, and W. W. Webb, 1974. Fluorescence correlation spectroscopy. II. An experimental realization. Biopolymers: Original Research on Biomolecules 13:29–61.

3. Magde, D., W. W. Webb, and E. L. Elson, 1978. Fluorescence correlation spectroscopy. III. Uniform translation and laminar flow. Biopolymers: Original Research on Biomolecules 17:361–376.

4. Denk, W., J. H. Strickler, and W. W. Webb, 1990. Two-photon laser scanning fluorescence microscopy. Science 248:73–76.

5. Schwille, P., U. Haupts, S. Maiti, and W. W. Webb, 1999. Molecular dynamics in living cells observed by fluorescence correlation spectroscopy with one-and two-photon excitation. Biophysical Journal 77:2251–2265.

6. Schwille, P., J. Korlach, and W. W. Webb, 1999. Fluorescence correlation spectroscopy with single-molecule sensitivity on cell and model membranes. Cytometry: The Journal of the International Society for Analytical Cytology 36:176–182.

7. Berland, K., P. So, and E. Gratton, 1995. Two-photon fluorescence correlation spectroscopy: method and application to the intracellular environment. Biophysical Journal 68:694–701.

8. Zernike, F., 1953. How I discovered phase contrast. Science 121:345–349.

9. Zernike, F., 1934. Beugungstheorie des Schneidenverfahrens und seiner verbesserten Form, der Phasenkontrastmethode. Physica 1:689–704.

10. Born, M., and E. Wolf, 1999. Principles of Optics: Electromagnetic Theory of Propagation, Interference and Diffraction of Light. Cambridge University Press.

11. Booth, M. J., 2014. Adaptive optical microscopy: the ongoing quest for a perfect image. Light: Science & Applications 3:e165.

12. Goodwin, E. P., and J. C. Wyant, 2006. Field Guide to Interferometric Optical Testing. SPIE Press, Bellingham, WA.

13. Lakshminarayanan, V., and A. Fleck, 2011. Zernike polynomials: a guide. Journal of Modern Optics 58:545–561.

14. Tyson, R., and V. Lakshminarayanan, 2012. Adaptive optics. Journal of Modern Optics 59:1032–1033.

15. Wikipedia, 2023. Zernike polynomials. https://en.wikipedia.org/wiki/Zernike_polynomials.

16. Enderlein, J., I. Gregor, D. Patra, and J. Fitter, 2004. Art and Artefacts of Fluorescence Correlation Spectroscopy. Current Pharmaceutical Biotechnology 5:155–161.

17. Enderlein, J., I. Gregor, D. Patra, T. Dertinger, and U. B. Kaupp, 2005. Performance of Fluorescence Correlation Spectroscopy for Measuring Diffusion and Concentration. ChemPhysChem 6:2324–2336.

18. Wolf, E., 1959. Electromagnetic diffraction in optical systems-I. An integral representation of the image field. Proceedings of the Royal Society of London. Series A. Mathematical and Physical Sciences 253:349–357.

19. B., R., W. E., and G. Dennis, 1959. Electromagnetic diffraction in optical systems, II. Structure of the image field in an aplanatic system. Proceedings of the Royal Society of London. Series A. Mathematical and Physical Sciences 253:358–379.

20. Fazel, M., K. S. Grussmayer, B. Ferdman, A. Radenovic, Y. Shechtman, J. Enderlein, and S. Pressé, 2024. Fluorescence microscopy: a statistics-optics perspective. Reviews of Modern Physics 96:025003.

21. Webb, W. W., 1976. Applications of fluorescence correlation spectroscopy. Quarterly Reviews of Biophysics 9:49–68.

22. Dertinger, T., V. Pacheco, I. von der Hocht, R. Hartmann, I. Gregor, and J. Enderlein, 2007. Two-focus fluorescence correlation spectroscopy: A new tool for accurate and absolute diffusion measurements. ChemPhysChem 8:433–443.

23. Müller, C. B., A. Loman, V. Pacheco, F. Koberling, D. Willbold, W. Richtering, and J. Enderlein, 2008. Precise measurement of diffusion by multi-color dual-focus fluorescence correlation spectroscopy. EPL (Europhysics Letters) 83:46001.

24. Petrášek, Z., and P. Schwille, 2008. Precise measurement of diffusion coefficients using scanning fluorescence correlation spectroscopy. Biophysical Journal 94:1437–1448.

25. Petrášek, Z., S. Derenko, and P. Schwille, 2011. Circular scanning fluorescence correlation spectroscopy on membranes. Optics Express 19:25006–25021.

26. Benda, A., M. Beneš, V. Marecek, A. Lhotsky, W. T. Hermens, and M. Hof, 2003. How to determine diffusion coefficients in planar phospholipid systems by confocal fluorescence correlation spectroscopy. Langmuir 19:4120–4126.

27. Humpolíčková, J., E. Gielen, A. Benda, V. Fagulova, J. Vercammen, M. Hof, M. Ameloot, Y. Engelborghs, et al., 2006. Probing diffusion laws within cellular membranes by Z-scan fluorescence correlation spectroscopy. Biophysical Journal 91:L23–L25.

28. Press, W. H., W. T. Vetterling, S. A. Teukolsky, and B. P. Flannery, 2007. Numerical Recipes 3rd Edition: The Art of Scientific Computing. Cambridge University Press.

